# LAG-3 blockade reactivates the CD8^+^ T cell expansion program to re-expand contracted clones in the tumor

**DOI:** 10.1101/2024.07.05.601623

**Authors:** Munetomo Takahashi, Mikiya Tsunoda, Hiroyasu Aoki, Masaki Kurosu, Haru Ogiwara, Shigeyuki Shichino, David Bending, Shumpei Ishikawa, James E.D. Thaventhiran, Kouji Matsushima, Satoshi Ueha

## Abstract

Effective cancer immunotherapy relies on the clonal proliferation and expansion of CD8^+^ T cells in the tumor^1,2^. However, our insights into clonal expansions are limited, owing to an inability to track the same clones in tumors over time. Here, we developed a multi-tumor mouse model system to track hundreds of expanding and contracting CD8^+^ T cell clones over multiple timepoints in tumors of the same individual. Through coupling of clonal expansion dynamics and single-cell RNA/TCR-seq data, we identified a transcriptomic signature in PD-1^+^Ly108^+^ precursor exhausted cells^3,4^ that strongly predicts rates of intratumoral clone expansion in mice and humans. We found that expression of the signature successfully stratifies melanoma patient outcomes to PD-1/PD-L1 blockade^5,6^. Downregulation of the signature precedes clone contraction – a phase in which clones contract but maintain revivable precursor exhausted cells in the tumor. LAG-3 blockade – an FDA-approved therapy whose effects on CD8^+^ T cell responses are currently unclear^7^, re-activates the expansion signature, re-expanding pre-existing clones, including previously contracted clones. These findings reveal how the study of clonal expansion dynamics provide a powerful ‘pan-immunotherapy’ signature for monitoring immunotherapies, including PD-1/PD-L1 and LAG-3 blockade, with implications for their future development.

## Introduction

Immunotherapies, centered around immune checkpoint blockade (ICB) therapies, have revolutionized cancer treatment by harnessing the power of our immune response^8^. Despite their success, challenges remain, particularly in overcoming treatment-resistant cancers^2^. While immunotherapies, including PD-1/PD-L1 and LAG-3 blockade, demonstrate clinical potential^1,7,9,10^, their development is hindered by our inability to comprehensively monitor their effect on T cell responses over time^2,9,11,12^, complicating efforts to accurately predict patient outcomes and optimize treatment protocols.

Recent landmark papers sampling tumors pre and post PD-1/PD-L1 blockade show that PD-1/PD-L1 blockade recruits novel T cell clones and revives pre-existing clones at the tumor site^13–16^, highlighting the need to track T cell responses in the same individuals to understand the effects of immunotherapy^2,11,12^. These studies have relied on longitudinally sampling the same patient tumors, limiting their broader application for investigating emerging immunotherapies, for which such patient samples are sparse^17^. Additionally, these studies only sample a small portion of the tumor, making it difficult to determine whether any observed changes reflect true biology, or are an artefact of the limited sampling^12^. While preclinical models have the potential to fill this gap – as demonstrated by a recent study time-stamping tumor-infiltrating CD8^+^ T cells^17^, they are also currently limited by their inability to comprehensively sample tumors over time.

Here, we have developed a multi-tumor mouse model system to track CD8^+^ T cell clones in tumors of the same individual over time. We inoculated mice with multiple tumors to overcome the challenges of sampling cells from the same tissue and utilized the T-cell receptor (TCR) sequence as a natural genetic barcode to identify clonally related T cell populations (“clones”) across tumors. By sequentially sampling tumors from the same mice, we successfully tracked hundreds of expanding and contracting clones over multiple timepoints in tumors of the same individual. We identified a transcriptomic CD8^+^ T cell signature that is highly predictive of clone expansion. This expansion signature successfully stratifies melanoma patient responses to PD-1/PD-L1 blockade. We observed that LAG-3 blockade upregulates the expansion signature, re-expanding pre-existing clones, including contracted clones. Together, these findings provide a time-resolved understanding of CD8^+^ T cell clonal responses that provide a powerful ‘pan-immunotherapy’ signature for monitoring immunotherapies, with implications for their future development.

## Results

### A time-resolved model of CD8^+^ T cell responses tracks expanding and contracting clones in the tumor

We, and others, have previously demonstrated that tumor growth and T cell infiltration is mirrored across bilaterally inoculated tumors^18–21^. We therefore investigated if a multi-tumor mouse model could be utilized to track T cell responses over time in the same individual. Lewis lung carcinoma (LLC) cells were inoculated into the left flank, right flank, and left hip of mice. The characteristics of tumor-infiltrating CD8^+^ T cell, as evaluated by PD-1, Ly108 and TIM-3 surface markers, were largely unaffected by the number of tumors implanted (Extended Data Fig. 1a-b). Across all tumors, including the left and right flank tumors, the sizes of any given clone (as identified by the read frequency of a TCR sequence in the repertoire) were equivalent (Fig. 1a-b, Extended Data Fig. 1c). These results indicate that for a given clone, each tumor acts as a proxy of clone size in the other tumors. We reasoned that comparison of clones across sequentially removed tumors could therefore enable the tracking of clones over time (Fig. 1c). LLC cells were bilaterally inoculated into the left and right flanks of mice (n=7). After 14 days, the left tumor was surgically removed, and a week later, the right tumor was harvested with the mice. CD8^+^ T cells obtained from the left and right tumor were analyzed by bulk TCR-sequencing (TCR-seq) to identify clones. Compared to clones obtained from tumors removed at the same time, clones obtained from sequentially removed tumors had greater variance in size, indicating ongoing expansion and contraction (Fig. 1d). Following established protocols^22–24^, we applied a beta-binomial model to define significantly expanded and contracted clones. Clones that were significantly larger in the tumor harvested at the later timepoint (right tumor) were defined “expanding clones” while clones that were significantly smaller were defined “contracting clones” (Fig. 1d, Extended Data Fig. 1d). Expanding and contracting clones were reflected by changes in cell count and identified in all mice (Fig. 1e-f, Extended Data Fig. 1e-f, indicating our system could reproducibly track clonal expansion dynamics.

**Fig. 1.**
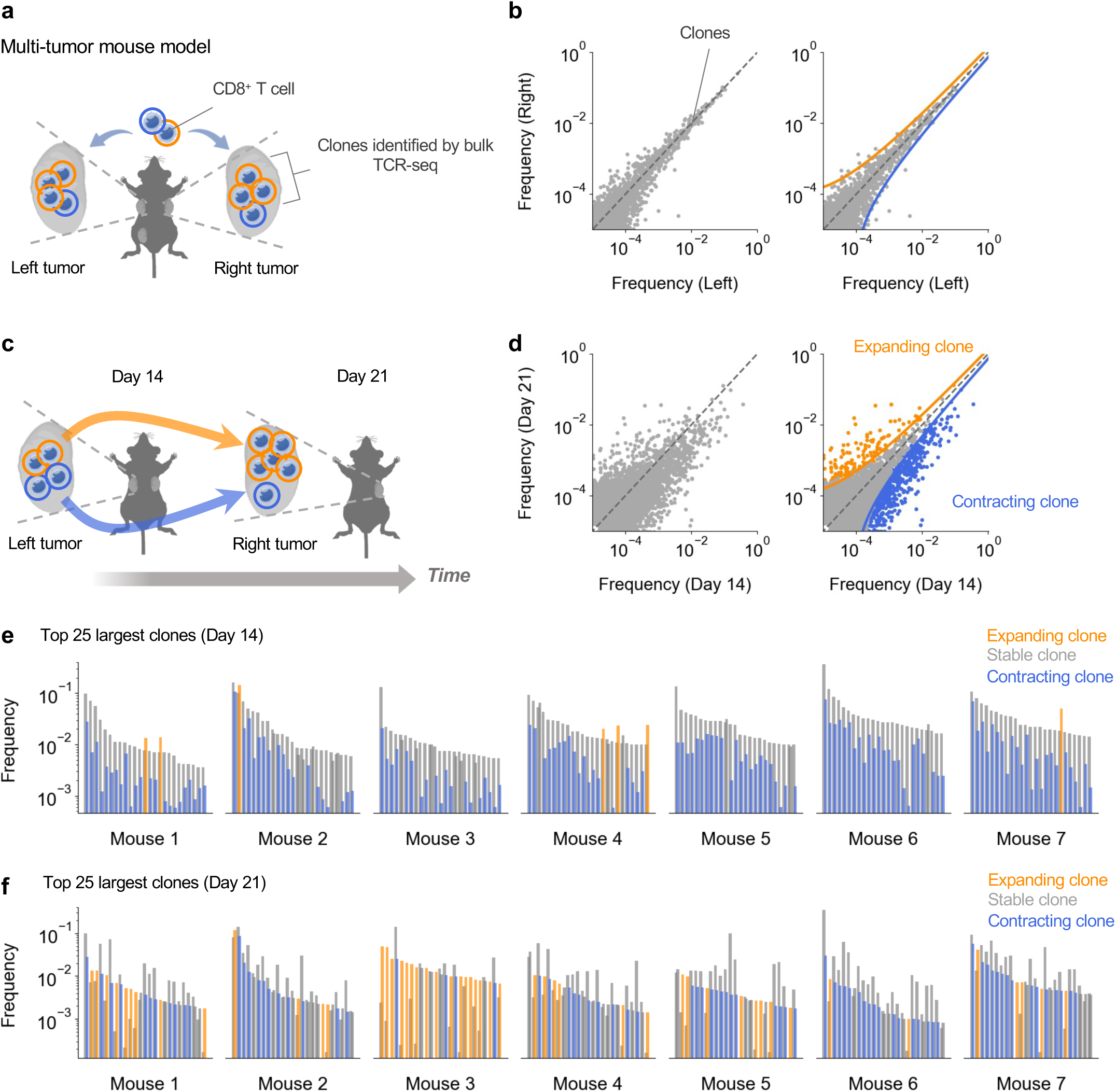
A time-resolved model of CD8^+^ T cell responses tracks expanding and contracting clones in the tumor. **a)** Depiction of a multi-tumor mouse model: Lewis lung carcinoma tumors are implanted simultaneously into the left flank, right flank and left hip of mice. Cells from the tumors are analyzed by bulk TCR-sequencing to identify cells belonging to the same clone. **b)** Scatter plots displaying the frequency of clones (normalized read count of each clone) in the left and right flank tumors of 3 mice excised at the same time (Left) with representative bounds defining expanding and contracting clones overlayed (Right). **c)** Bilateral tumors were excised independently on sequential days in the same mice to track the temporal dynamics of clones in the tumor. **d)** Scatter plots displaying the frequency of clones in tumors on days 14 and 21 from 7 mice (Left) with representative bounds defining expanding and contracting clones overlayed (Right). Clones colored by their expansion dynamics as defined using a beta-binomial model from Rytlewski et al., 2019^22^. **e-f)** The frequency of the top 25 largest clones obtained from each mouse on day 14 **(e)** and 21 **(f)** overlayed with the frequency of the same clone on day 21 **(e)** and 14 **(f)** respectively. In both figures, clone frequencies on day 21 are colored by their expansion dynamics. Dots represent clones (**b, d**).

### Precursor exhausted cells drive clone expansion and are preferentially maintained in the tumor during clone contraction

To characterize the differentiation states of expanding and contracting clones, we sorted intratumoral CD8^+^ T cells based on PD-1, Ly108 and TIM-3 expression before processing them through bulk TCR-seq (Fig. 2a, Supplementary Table 1). This analysis identified the proportion of cells in each clone that were PD-1^−^, precursor exhausted (PD-1^+^Ly108^+^TIM-3^−^) – a stem cell like state capable of self-renewal^3,4^, intermediate exhausted (PD-1^+^Ly108^+^TIM-3^+^) and terminally exhausted (PD-1^+^Ly108^−^TIM-3^+^) – a short-lived terminally differentiated state^3^ on days 14 and 21. Following accumulating evidence that terminally exhausted cells in tumors are specifically derived from tumor-specific T cells^14,25,26^, we followed published protocols and filtered for clones that contained counts in the terminally exhausted compartment to enrich for tumor reactive clones^14^ (Extended Data Fig. 2a). Comparison of the exhaustion states of clones on day 14 revealed that expanding clones contained a higher fraction of precursor exhausted cells, and a lower fraction of terminally exhausted cells than contracting clones (Extended Data Fig. 2b-c). We tracked the changes in exhaustion states over time (Fig. 2b). Expanding clones decreased their fraction of precursor exhausted cells and increased their fraction of terminally exhausted cells, consistent with the differentiation of precursor exhausted cells into terminally exhausted cells^3^. In contrast, contracting clones, particularly those with high fractions of terminally exhausted cells ‘regressed’ – they decreased their fraction of terminally exhausted cells and increased their fraction of precursor exhausted cells. We analyzed changes in the number of precursor exhausted and terminally exhausted cells in each clone (Fig. 2c). For many clones with high fractions of terminally exhausted cells, the number of terminally exhausted cells decreased while the number of precursor exhausted cells decreased less, or in some cases, increased (Fig. 2d). Collectively, these results indicate that precursor exhausted cells differentiate to drive clone expansion and are preferentially maintained in the tumor during clone contraction (Fig. 2e).

**Fig. 2.**
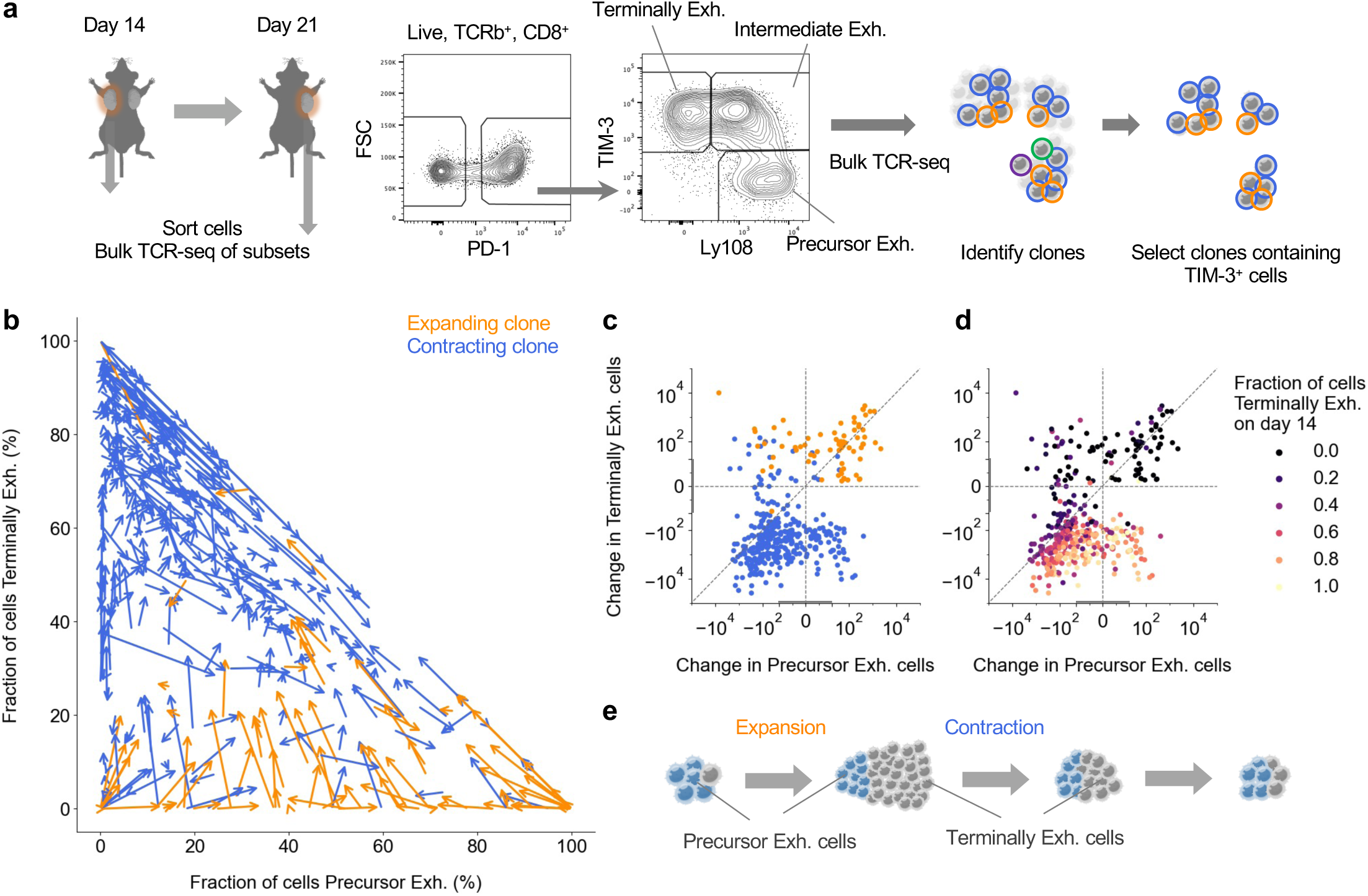
Precursor exhausted cells drive clone expansion and are preferentially maintained in the tumor during clone contraction. **a)** Cells from tumors excised on days 14 and 21 (n=7 mice) were sorted based on marker expression before processing through bulk TCR-sequencing, enabling phenotypic characterization of individual clones. Clones containing TIM-3^+^ cells were selected for downstream analysis as in Extended Data Fig. 2a. **b)** Velocity plot depicting the cellular composition changes of expanding and contracting clones from day 14 to 21. Arrows depict individual clones. The base of the arrow indicates the fraction of cells that were precursor exhausted and terminally exhausted on day 14 while the arrow direction and length indicate the magnitude and vector of change in these fractions over time. **c-d)** Change in the number of precursor and terminally exhausted cells between day 14 and 21 colored by their expansion dynamics (**c**) and fraction of cells terminally exhausted on day 14 (**d**). Symmetrical log axis with a linear range of –20 to +20 (as depicted by grey lines on axes). **e**) Expansion and contraction phases of clones. Expanding and contracting clones from the 7 mice that displayed changes in both the precursor exhausted and terminally exhausted cell fractions are shown (**b-d**). Dots represent clones (**c, d**).

### A transcriptomic signature in precursor exhausted cells predicts clone expansion

Since both expanding and contracting clones contained similar numbers of precursor exhausted cells per clone (Extended Data Fig. 2d), we hypothesized that differences in their expansion dynamics could be attributed to the state of their precursor exhausted cells. We utilized the same bilateral tumor excision schema and performed single-cell RNA/TCR-seq on a fraction of T cells, enriched for precursor exhausted cells, in addition to bulk TCR-seq (Extended Data Fig. 3a, Supplementary Table 1). This produced a time-resolved dataset that annotated single CD8^+^ T cells with their transcriptomic states and their clonal expansion dynamics. Unsupervised clustering of the single-cell RNA/TCR-seq data retrieved precursor exhausted (clusters 0, 1 and 4), intermediate exhausted (clusters 2, 5 and 7) and terminally exhausted cells (cluster 3) (Fig. 3a, Extended Data Fig. 3b-c). We compared the transcriptomic state of precursor exhausted cells from expanding and contracting clones on day 14 to find markers that would predict their future clonal expansion dynamics (Fig. 3b). Precursor exhausted cells from expanding and contracting clones were mainly enriched in a precursor exhausted cluster (cluster 0) that highly expressed *Lag3* and lowly expressed *Sell* (Fig. 3c, Extended Data Fig. 3c). These cells could not be differentiated based on their Uniform Manifold Approximation and Projection (UMAP) representation (Fig. 3c, Extended Data Fig. 3d). However, differential gene expression (DEG) analysis between precursor exhausted cells from expanding and contracting clones revealed that cells from expanding clones overexpressed 45 genes – of which 22 were canonical genes (and not mitochondrial or ribosomal genes) which we termed the ‘expansion’ signature (Fig. 3d, Supplementary Tables 2 and 3). The expansion signature contained genes associated with T cell memory states (*Tcf7*, *Ccr7*), T cell signaling (*Ptprcap*, *Cd160*), and cell cycle regulation (*Hmgn1, Ddit4, Rack1*). As expected, precursor exhausted cells from clones that later expanded upregulated the expansion signature while cells from clones that later contracted downregulated the signature (Extended Data Fig. 3e). We tested if, in addition to predicting expansion or contraction of the whole clone, expression of the expansion signature could also predict the extent of clone expansion (the log2 fold-increase in clone frequency from day 14 to 21). Strikingly, the expansion signature score correlated with clone expansion (Fig. 3e). The correlation was stronger when we scored the expansion signature of individual clones by averaging the score of their constituent precursor exhausted cells (Fig. 3f). The correlations were evident when we analyzed each mouse independently (Extended Data Fig. 3f) and altered the analysis criteria (Extended Data Fig. 3g), Other previously reported precursor exhausted cell state gene signatures from mice and humans did not consistently correlate with the expansion of clones (Extended Data Fig. 3h).

**Fig. 3.**
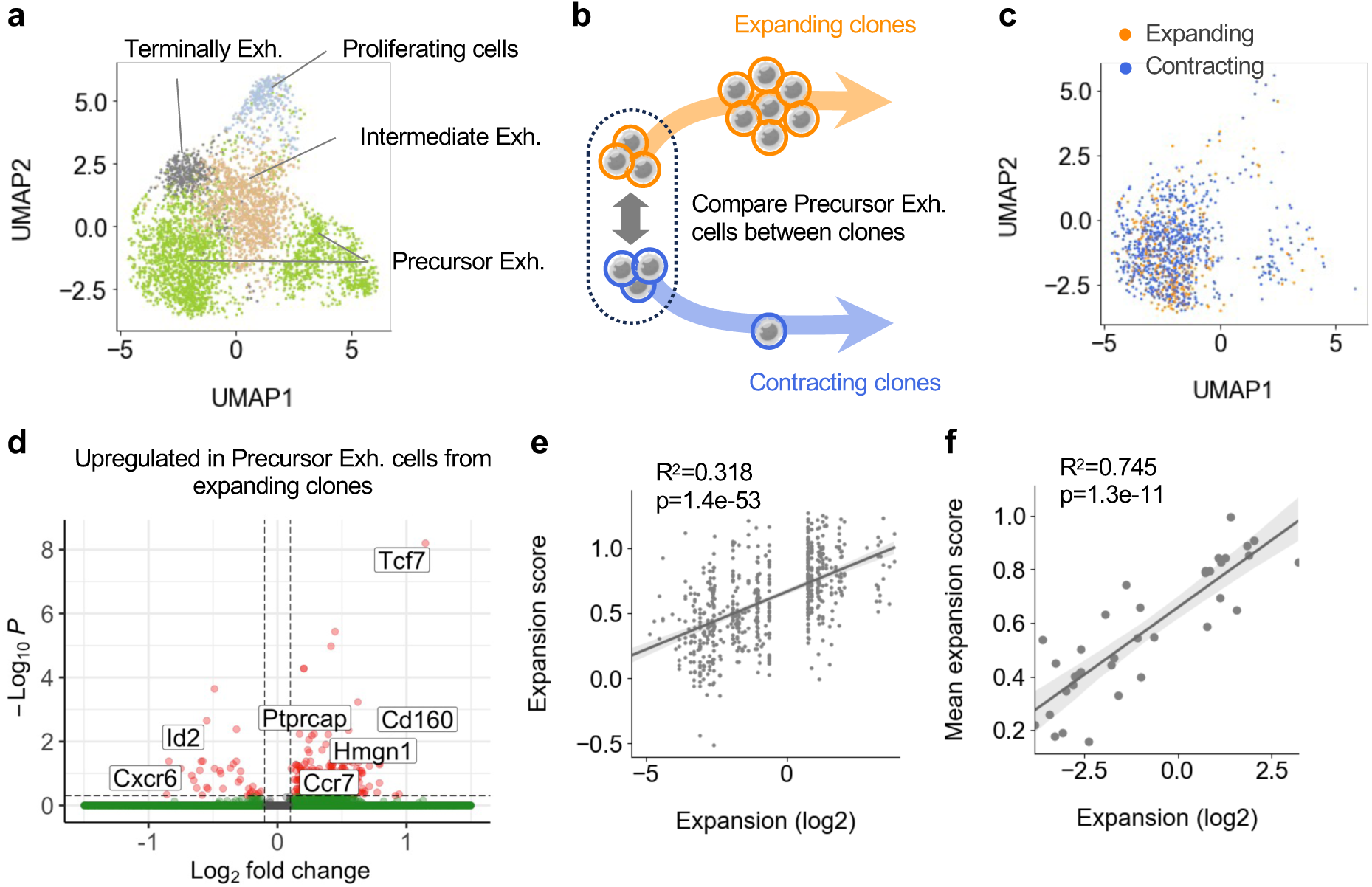
A transcriptomic signature in precursor exhausted cells predicts clone expansion. Cells from bilateral tumors excised on days 14 and 21 from 3 mice were processed by a combination of single-cell RNA/TCR-sequencing and bulk TCR-sequencing to enable integration of single-cell transcriptomic data of clones with their expansion dynamics. **a)** Uniform Manifold Approximation and Projection (UMAP) map of 4461 cells from tumors excised on day 14, colored and labelled by cell type. **b-f)** Precursor exhausted cells from tumors excised on day 14, belonging to expanding and contracting clones, were investigated for transcriptomic signatures that could predict future clone expansion. **c)** UMAP map of precursor exhausted cells colored by their expansion dynamics. **d**) Differential gene expression analysis of genes overexpressed in precursor exhausted cells from expanding clones over contracting clones. Notable genes are labelled. **e**) Scatter plot comparing the expansion signature score with the expansion (the log2 fold-increase in clone frequency from day 14 to 21) of the clone for individual cells. Cells belonging to the same clone have the same expansion rate. **f**) Scatter plot comparing the mean expansion signature score with the expansion of clones. Clones were scored for their expression of the expansion signature in constituent precursor exhausted cells on day 14. Analysis on clones with at least 5 precursor exhausted cells on day 14. Dots represent genes (**d**), cells (**a**, **c, e**) and clones (**f**). Statistical testing via Wilcoxon rank-sum test corrected with the Benjamini–Hochberg procedure (**d**) and Wald test (**e, f**) (****, p<0.0001; ***, p<0.001; **, p<0.01; *, p<0.05; ns p>0.05).

To undertake an unbiased assessment of the pathways associated with the expansion signature, we scored precursor exhausted cells of clones for hallmark pathway gene signatures^27^ – 50 gene signatures of well-defined biological processes, and searched for correlations with the expansion signature score at the clonal level. Hallmark pathways associated with increased metabolism (oxidative phosphorylation, glycolysis, adipogenesis, fatty acid metabolism) and proliferation (MYC targets, MTORC1 signaling) positively correlated with the expansion signature score, whereas a pathway for TGF-B signaling inversely correlated with the expansion signature score (Extended Data Fig. 4a). Recent studies have implicated a role for TGF-B and Let-7 signaling in repressing cellular metabolism through inhibition of MYC signaling^28–30^. In our dataset, clones with low expansion signature scores highly expressed genes associated with TGF-B signaling in T cells^29,31^ whereas clones with high expansion signature scores highly expressed genes inhibited by Let-7^30^ (Extended Data Fig. 4c). These findings were replicated in precursor exhausted cells from a single-cell RNA/TCR-seq dataset of antigen-signaled CD8^+^ T cells infiltrating the YUMMER1.7 model^32^ (Extended Data Fig. 4b, d). Taken together, these results demonstrate the identification of an expansion signature, associated with high metabolic activity and reduced TGF-B signaling, that is highly predictive of the extent of future clone expansions.

### The expansion signature stratifies melanoma patient outcomes to PD-1 and PD-1/CTLA-4 blockade

We explored whether the expansion signature might be applicable to humans. We analyzed a single-cell RNA/TCR-seq dataset obtained from a longitudinal study of basal cell carcinoma pre and post PD-1/PD-L1 blockade from Yost et al., 2019^13^. While expression of the expansion signature did not correlate with expansion at the cellular level, a strong correlation was observed when cells were grouped into clones (Fig. 4a, Extended Data Fig. 5a-b). Comparing responders and non-responders, clones from responders in post-treatment samples had significantly higher expansion signature scores than clones from non-responders, while no such difference was observed in pre-treatment samples (Fig. 4b, Extended Data Fig. 5c). These results suggested that response is associated with high clonal expansion activity, leading us to query whether the expansion signature could be used to monitor immunotherapy outcomes. We scored bulk RNA-seq melanoma tumor biopsies of pre– and on-PD-1/PD-L1 blockade treatment patients from Riaz et al., 2017^5^ with the expansion signature score. We utilized a stringent expansion gene signature (adjusted p-value cut-off of 0.01 instead of 0.05, Supplementary Table 2) to account for the increased biological variability in tumor biopsies. Corroborating our previous analysis, on-treatment (but not pre-treatment) biopsies from responders scored significantly higher for the expansion signature than non-responders (Extended Data Fig. 5d). Patients in this dataset were either treatment naive or were treated with CTLA-4 blockade prior to PD-1/PD-L1 blockade treatment.

**Fig. 4.**
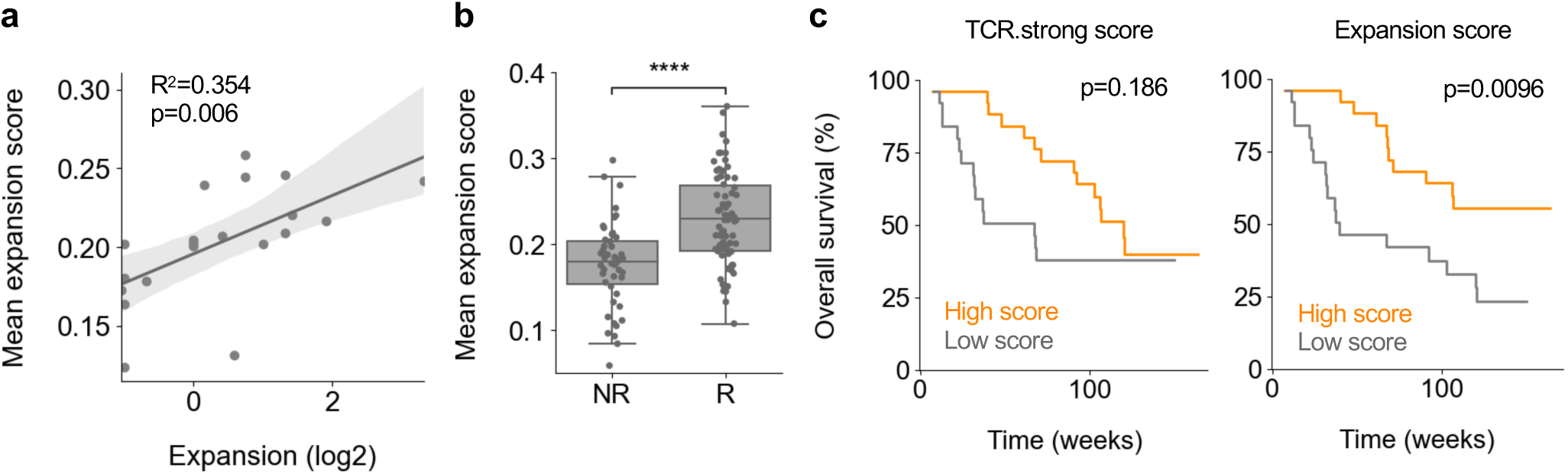
The expansion signature stratifies melanoma patient outcomes to PD-1/PD-L1 blockade. **a-b**) Analysis of CD8^+^ T cells from site-matched primary tumor single-cell RNA/TCR-seq samples from advanced basal cell carcinoma patients pre and post anti-PD-1 treatment from Yost et al., 2019^13^. **a)** Scatter plot showing correlation between the mean expansion signature score in cells of clones pre-treatment and the expansion of the clone (the log2 fold-increase of clone count from pre-to post-treatment). Analysis of clones with at least 2 cells pre-treatment. **b)** Comparison of mean expansion score in cells of clones post-treatment between non-responders (NR) and responders (R). Analysis of clones with at least 2 cells post-treatment. **c)** Analysis of bulk RNA-seq tumor biopsies from melanoma patients (n=50) during anti-PD-1 treatment from Riaz et al., 2017^5^. Plots of Kaplan Meier estimator of overall survival based on median of TCR.strong (Left) or expansion (Right) signature scores. Dots represent clones (**a-b**). Statistical testing via Wald test (**a**), Kruskal-Wallis test (**b**) and Log-rank test (**c**) (****, p<0.0001; ***, p<0.001; **, p<0.01; *, p<0.05; ns p>0.05).

Elliot et al., 2021^33^ recently reported a TCR.strong metric (consisting of *TNFRSF4*, *IRF8*, *STAT4*, *TNIP3* and *ICOS* derived from strong TCR signaling) that could stratify patient outcomes for treatment naïve patients in this dataset. Following their protocols^33^, we split patients into a “High” or “Low” cohort based on the median expansion signature score and (for comparison) TCR.strong score and plotted corresponding survival curves. Strikingly, the “High” expansion signature score cohort showed a significant increase in overall survival compared to the “Low” cohort without splitting the cohort based on previous exposure to CTLA-4 blockade, whereas no such difference was seen with the TCR.strong score (Fig. 4c). The “High” expansion signature score cohort demonstrated better overall survival when each treatment regimen was analyzed separately (Extended Data Fig. 5e), indicating the power of the expansion signature in stratifying different treatment regimen outcomes. In concordance with overall survival, the expansion signature score similarly performed better than the TCR.strong score for stratifying progression free survival (Extended Data Fig. 5f). We validated that the expansion signature could stratify immunotherapy outcomes in a separate dataset of early-during-treatment (EDT) patients undergoing PD-1/PD-L1 or PD-1/PD-L1 and CTLA-4 blockade from Gide et al., 2019^6^ (Extended Data Fig. 5g-h). Together, these results indicate the immediate utility of the expansion signature as a powerful ‘pan-immunotherapy’ signature for monitoring immunotherapy outcomes.

### LAG-3 blockade enhances the expansion signature and re-expands contracting clones in the tumor

To investigate how the expansion signature changes in clones over time, we analyzed our time-resolved day 21 single-cell RNA/TCR-seq dataset (Extended Data Fig. 6a, Supplementary Table 1), focusing on precursor exhausted cells from expanding and contracting clones. Compared to terminally exhausted cells, precursor exhausted cells from both expanding and contracting clones continued to express genes associated with their stem-cell like state on day 21 (Extended Data Fig. 6b). From day 14 to 21, cells from expanding clones decreased their expansion signature scores (Extended Data Fig. 6c). Conversely, cells from contracting clones increased their expansion signature scores (Extended Data Fig. 6c). These results, combined with our finding that precursor exhausted cells from contracting clones are preferentially maintained in the tumor, indicate a possibility to re-expand contracted clones by reactivation of their precursor exhausted cells.

Following the development of CTLA4 and PD-1/PD-L1 blockade, Lymphocyte activation gene-3 (LAG-3) blockade has emerged as the third checkpoint inhibitor to be approved for clinical use^7^. While the mechanisms by which it enhances the T cell response remains incompletely understood, there is evidence suggesting it could reduce TGF-B signaling^7,34^. We therefore investigated whether LAG-3 blockade could re-activate the expansion signature in precursor exhausted cells. Tumors were excised 4 days after LAG-3 blockade, and the precursor exhausted cells (subset for CD62L^−^ to enrich for expanding and contracting clones – cluster 0 in Extended Data Fig. 3c) was analyzed by bulk RNA-seq (Fig. 5a). Geneset enrichment analysis (GSEA) revealed that LAG-3 blockade significantly upregulated their expansion signature, and significantly downregulated their TGF-B signaling signature (Fig. 5b-c), indicating that LAG-3 blockade was reactivating the clone expansion signature.

**Fig. 5.**
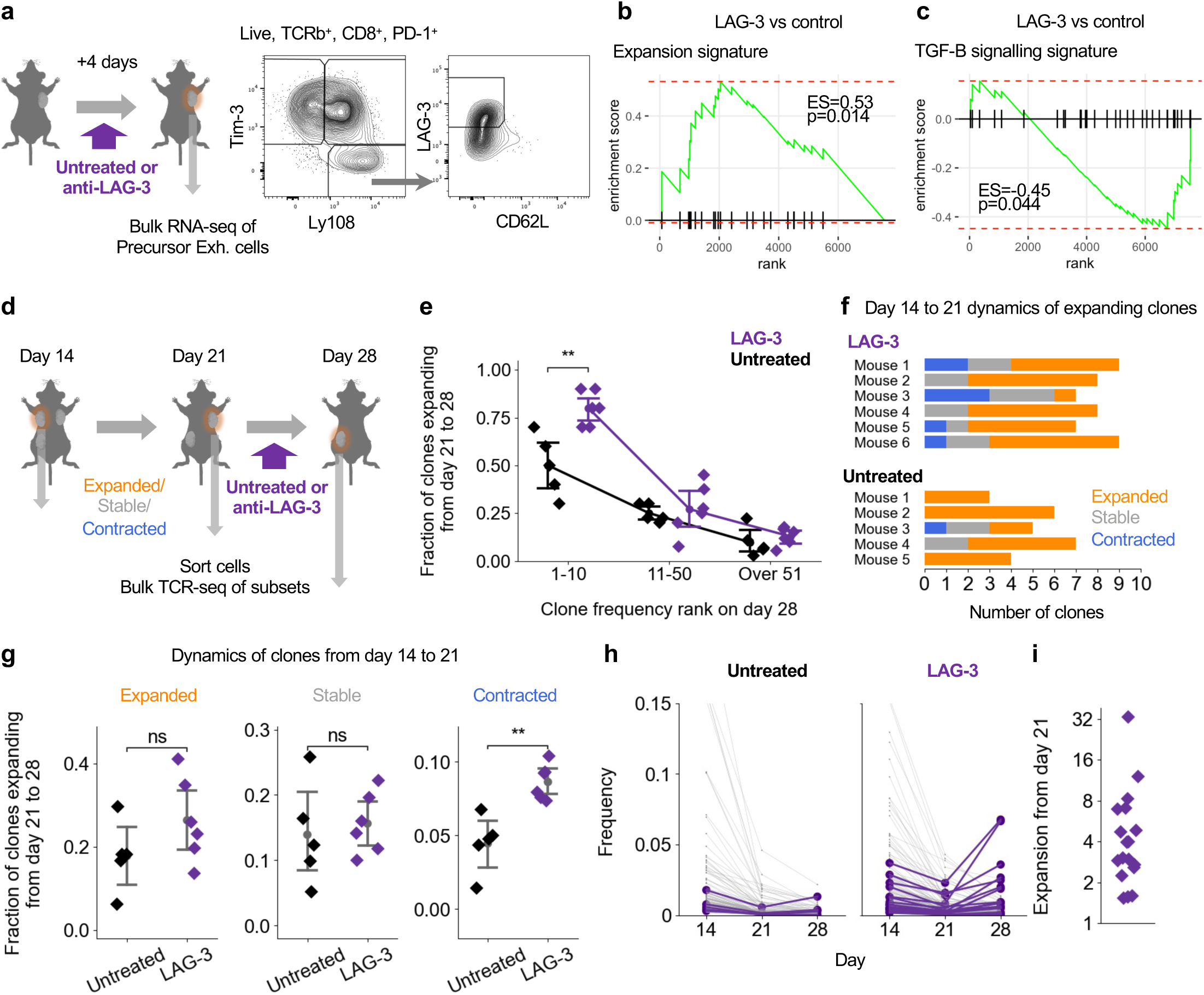
LAG-3 blockade enhances the expansion signature and re-expands contracting clones in the tumor. **a)** Tumors were resected from mice on day 18, 4 days after starting anti-LAG-3 or no treatment (n=6, 5 for anti-LAG-3 and no treatment respectively). **b-c)** Geneset enrichment analysis (GSEA) testing enrichment of the expansion signature (**b**) and TGF-B signaling signature **(c)** in genes overexpressed in the precursor exhausted cells after anti-LAG3 treatment over untreated mice. Enrichment score (ES) and p values as shown. **d)** Tumors were resected sequentially from multi-tumor mice as shown. After day 21, mice were treated with anti-LAG-3 or no treatment, (n=6, 5 for anti-LAG-3 and no treatment respectively). Clones containing TIM-3^+^ cells were selected for downstream analysis. **e**) Fraction of clones detected in the tumor on day 28 expanding between day 21 and 28, grouped by the ranked size of the clone on day 28 and colored by each treatment regimen. **f**) Expanding clones from the top 10 clones at day 28 (as compared in **e**) colored by their day 14 to 21 expansion dynamics. **g**) Fraction of clones expanding between day 21 to 28 for clones that expanded (Left), were stable (Middle) or contracted (Right) between day 14 and 21. **h**) The change in frequency between day 14 and 28 of the largest clones (showing top 50 clones on day 14 or 21) that contracted between day 14 and 21. Clones expanding between day 21 and 28 depicted in purple. **i**) Expansion (the log2 fold-increase in clone frequency from day 21 to 28) of clones from (**h**) after anti-LAG-3 treatment. Dots represent clones (**h, i**) and mice (**e, g**) with mean and 95% confidence interval as shown (**e, g**). Statistical testing via Kruskal-Wallis test (**e, g**) (****, p<0.0001; ***, p<0.001; **, p<0.01; *, p<0.05; ns p>0.05).

We employed our multi-tumor model to assess if LAG-3 blockade did indeed increase CD8^+^ T cell clone expansions. Tumors were inoculated into the left flank, right flank and left hip of mice and sequentially removed on days 14, 21 and 28 to enable the tracking of clones over three time-points. Mice were administered LAG-3 blocking antibodies or left untreated after removal of their second tumor on day 21 (Fig. 5d). This experimental schema allowed us to assess the effect of LAG-3 blockade on clonal expansion dynamics and differentiate its effect between previously expanded and contracted clones. We analyzed clones that were newly detected on day 28, and clones that had persisted from previous timepoints. Contrary to the reported effects of PD-1/PD-L1 blockade^13–15^, LAG-3 blockade did not significantly increase the fraction of newly detected clones in the tumor (Extended Data Fig. 7a). Rather, LAG-3 blockade significantly increased the fraction of large clones that were expanding (Fig. 5e). We observed a consistent trend in the fraction of large clones that were contracting (Extended Data Fig. 7b). We labelled each of the large, expanding clones by their day 14 to 21 dynamics (Fig. 5f). Compared to the untreated control, many of the clones that were expanding after LAG-3 blockade had contracted between day 14 and 21. When we analyzed all clones, we observed that LAG-3 blockade significantly increased the fraction of contracted clones that were (re-)expanding by almost two-fold (Fig. 5g). Several contracted clones were re-expanding to frequencies more than 5 times higher than their day 21 frequencies – with one clone re-expanding 33-fold (Fig. 5h-i). These expansions altered the characteristics of CD8^+^ T cells infiltrating the tumor. LAG-3 blockade increased the combined frequency of expanding clones by almost 20% (from 36.3% to 55.6%, on average) (Extended Data Fig. 7c) with most of this increase attributable to expanding clones that were stable or had contracted between day 14 and 21 (Extended Data Fig. 7d). Together, these results demonstrate that LAG-3 blockade enhances the expansion signature and re-expands contracted clones in the tumor.

## Discussion

In this study, we utilized the multi-tumor mouse model to longitudinally assess CD8^+^ T cell responses to tumors. We identified a transcriptomic gene signature that predicts the expansion of CD8^+^ T cell clones. We validated this signature in other mice and human datasets and observed that LAG-3 blockade upregulates the signature and re-expands contracted clones. Our study of the effects of LAG-3 blockade provide the rationale for its potency in combination therapy^7^ – with PD-1/PD-L1 blockade effectively recruiting novel clones into the tumor site^13–15^, LAG-3 blockade assists by re-expanding clones already existent in the tumor. Application of the model to understand the cellular effects of other emerging immunotherapies could similarly provide insights required for optimizing their efficacy.

In our study, we demonstrated that the expansion signature stratifies outcomes for melanoma patients undergoing different immunotherapy regimens. Coupled with our finding that LAG-3 also upregulates the expansion signature, our results demonstrate the utility of the signature in all FDA-approved immune checkpoint blockade inhibitors. Given that almost all immunotherapies, including CAR/TCR-T cell therapies, vaccines, and monoclonal antibodies, rely on CD8^+^ T cell expansions for their success, we envision the expansion signature could form the basis for an even more generic ‘pan-immunotherapy’ signature that enables accurate monitoring of patient outcomes for existing treatments, and cost-effective evaluation of immunotherapies in development.

Our results also indicate the potential of targeting the expansion signature for developing potent immunotherapies. One of the transcription factors in the expansion signature, *Tcf7*, is well-characterized as regulating the development of the precursor exhausted state^35^ and serves as a biomarker for responders to immune checkpoint blockade therapies in clinical settings^36^. Our work suggests that *Tcf7* also plays a role in regulating the expansion of clones. Targeted alterations of *Tcf7* expression, and expression of the other genes identified in the expansion signature could enable regulation of clonal expansion dynamics. These interventions would be particularly transformative for CAR/TCR-T therapy.

More broadly, our results indicate that future clone expansion characteristics are, at least in part, encoded in transcriptomic profiles. This leaves exciting questions to pursue – could transcriptomic profiles of CD8^+^ T cells encode other temporal characteristics, such as differentiation fate? Is a large component of temporal CD8^+^ T cell behaviors pre-determined? We posit that similar approaches combining multimodal snapshot analyses with temporal tracking could identify transcriptomic signatures relevant for other clonal expansion dynamics, including the expansion of engrafted stem cells and the development of malignant tumors.

## Methods

### Experimental Mice and Cell Lines

All animal experiments were conducted in accordance with institutional guidelines with the approval of the Animal Care and Use Committee of Tokyo University of Science. Eight-week-old female C57BL/6 mice (CD90.2, RRID: MGI:5488963) were purchased from Sankyo Labo Service Corporation Inc. All mice were bred at specific pathogen-free facilities at Tokyo University of Science. The Lewis lung carcinoma (LLC) cell line was provided by Nihonkayaku (Tokyo, Japan).

### Tumor Model and Treatment

LLC cells (5 × 10^5^ cells) were inoculated subcutaneously into the left flank, right flank and (for experiments with three tumors) the left hip of C57BL/6J mice. In all experiments, except for control experiments to validate the proportional infiltration of clones, tumors were excised sequentially in the order of left flank tumor, right flank tumor and (for experiments with three tumors) left hip on days 14, 21 and (for experiments with three tumors) 28 or days 14 and 18 (for the bulk RNA-seq experiment). Tumor diameter was measured twice weekly and used to calculate tumor volume (mm^3^) [(major axis; mm) × (minor axis; mm)^2^]. Tumors were excised by sedating and opening a slit in the skin of the mouse, detaching the tumor from the peritoneum and the skin along with surrounding fat and draining lymph node, and clipping the slit back together. For treatment, monoclonal anti–PD-L1 (clone 10F.9G2, BioLegend) and anti-LAG-3 (clone C9B7W, BioLegend or Selleck) was injected intraperitoneally at a dose of 200 μg per mouse on days 14 and 16 (for bulk RNA-seq data) or days 21 and 23 (for bulk TCR-seq data) after tumor inoculation. Mice with incomplete excision resulting in immediate (<7 days) regrowth of excised tumors, or tumors that did not grow (<100mm^3^ by day 28) were excluded from the analysis.

### Generation of Single-Cell Suspensions from Tumors

Tumors were cut into small fragments and digested for 45 minutes at 37°C with 0.2% collagenase (FUJIFILM Wako) and DNase I (Sigma-Aldrich), except for experiments for single-cell analysis, in which tumors were digested with 0.2% collagenase NB8 (Nordmark Pharma GmbH). The cells were then subjected to two-phase density separation with upper 35% Percoll PLUS (GE Healthcare) (1000g, 10 minutes at room temperature). Leukocytes were recovered, washed and resuspended in preparation medium (PM) [5%FBS, 10mM HEPES (Nacalai), Penicillin-Streptomycin (Nacalai) in RPMI-1640 (Nacalai)]. After RBC lysis with ACK buffer [155mM of NH_4_Cl (Nacalai), 10mM of KHCO_3_ (Nacalai), and 0.5mM EDTA pH8.0 (NIPPON GENE)] for 1 minute at room temperature, cells were washed and resuspended in PM, and the cell concentrations were measured using Flow-Count fluorospheres (Beckman Coulter) and a CytoFLEX S flow cytometer (Beckman Coulter).

### Cell Staining and Pre-Enrichment

Cells were stained with a mix of Fc Block (anti-mouse CD16/CD32 mAb; clone 2.4G2, BioLegend) and fluorophore-conjugated anti-mouse monoclonal antibodies (Supplementary Table 5). Antibody mixtures were prepared using BD Horizon Brilliant Stain Buffer Plus (BD) diluted 5x with FACS buffer (D-PBS(-) supplemented with 2% FBS, and 0.05% Sodium Azide). Staining was performed for 20 minutes at 4°C. In the single-cell RNA-seq/TCR-seq experiment, cells were stained with antibodies and anti-CD45 Sample Tag oligonucleotide-conjugated antibodies from the Single-Cell Multiplexing Kit (BD Biosciences). Before sorting, T cells were enriched from tumor suspensions by further staining with BD IMag APC Magnetic Particles-DM (BD Biosciences), then placed on magnetic plate. After washing by MACS buffer (D-PBS(-) supplemented with 10% BSA, and 2 mM EDTA pH8.0), APC and APC-Cy7(CD8, mouse TCRβ)-positive cells were collected as positive fraction.

### Cell Sorting

Cell sorting was performed using a FACS Aria II or Aria III (BD Biosciences). Cells were stained with 5µg/mL of propidium iodide (PI, Sigma-Aldrich) immediately before sorting. Nonviable cells were excluded from the analysis based on forward and side scatter profiles and PI staining. For bulk TCR repertoire analysis and bulk RNA-seq analysis, cells were sorted into Lysis buffer [1% w/v LiDS, 100 mM Tris-HCl pH7.5, 500 mM LiCl, 5 mM dithiothreitol, and 10 mM EDTA pH8.0] and stored at −80°C. For single-cell RNA/TCR-seq, cells were sorted into D-PBS(-) with 10% FBS.

### Sample Collection

For the bulk TCR-seq experiments, cells were sorted from the CD8^+^, CD8^+^PD-1^+^Ly108^+^TIM-3^−^, CD8^+^PD-1^+^Ly108^+^TIM-3^+^, CD8^+^PD-1^+^Ly108^−^TIM-3^+^ and CD8^+^PD-1^−^ fractions to produce five datasets per mouse. For the single-cell RNA/TCR-seq and bulk TCR-seq experiment, cells were sorted from the CD8^+^PD-1^+^Ly108^+^TIM-3^−^, CD8^+^PD-1^+^Ly108^+^TIM-3^+^, CD8^+^PD-1^+^Ly108^−^TIM-3^+^ and CD8^+^PD-1^−^ fractions. From each of the fractions, a fixed number of cells was aliquoted for single cell sequencing, and the rest was processed by bulk TCR-seq. The numbers of cells collected for each data in each analysis are shown in Supplementary Table 1. For bulk RNA-seq samples, cells were sorted from the CD8^+^PD-1^+^Ly108^+^TIM-3^−^CD62L^−^ fraction.

### Library Preparation and Sequencing for TCR Repertoire Analysis

TCR sequencing libraries for next-generation sequencing were prepared as previously reported with two modifications^19^. In the first and second amplifications, 18 and 10 cycles of polymerase chain reaction (PCR) were performed respectively. Oligonucleotide sequences used in this study are shown in Supplementary Table 4.

### Library Preparation and Sequencing for Single Cell Analysis

Single cell sequencing libraries were prepared as previously reported^15^. Briefly, live cells were stained with Calcein AM (Nacalai) and 20,564 cells were loaded on an BD Rhapsody cartridge (BD) and processed using BD Rhapsody Cartridge Reagent Kit (BD) following the manufacturer’s instructions. Single-cell cDNA synthesis was performed using BD Rhapsody cDNA Kit (BD) following the manufacturer’s protocol. Whole transcriptome analysis (WTA), TCRseq, and Sample tag Index libraries were prepared using Immune Response Panel Mm (BD) according to the manufacturer’s instructions with modifications^15^. Amplified libraries were quantified using a KAPA SYBR Fast qPCR Kit (KAPA Biosystems) and its size distribution was analyzed by a MultiNA. WTA, TCRseq, and Sample tag libraries were sequenced on an Illumina Novaseq 6000 S4 flowcell (67 bp read 1 and 140 bp read 2) using NovaSeq 6000 S4 Reagent Kit v1.5 to a depth of approximately 20,000 reads, 7,000 reads, and 10,000 reads per cell, respectively.

### Library Preparation and Sequencing for Bulk RNA Sequencing Analysis

Bulk RNA-seq library preparation was performed with on-beads reverse transcription and template-switching. mRNA-trapped oligo-dT-immobilized Dynabeads M270-streptavidin (Thermo Fisher Scientific) were washed as in bulk TCR-seq library preparation, and resuspended in 10 µL of RT mix [1× First Strand buffer (Thermo Fisher Scientific), 1 mM dNTP, 2.5 mM DTT (Thermo Fisher Scientific), 1 M betaine (Sigma-Aldrich), 9 mM MgCl_2_ (NIPPON GENE), 1 U/µL RNaseIn Plus RNase Inhibitor (Promega, Madison, WI), 10 U/µL Superscript II (Thermo Fisher Scientific), and 1 µM of trP1-TSO], and incubated for 60 min at 42°C and immediately cooled on ice. Beads were washed once with B&W-T buffer [5 mM Tris-HCl pH 7.5, 1 M NaCl (NACALAI TESQUE), 0.5 mM EDTA, and 0.1% Tween-20 (Sigma-Aldrich)], and once with 10mM Tris-HCl pH 8.0. To amplify the WTA library, cDNA-immobilized beads were resuspended with 25 µL of the first PCR mixture [0.4 μM of primers (illumina-i7-BCXX-trP1, and 5’ BDWTA V2 primer), and 1x KAPA Hifi Hotstart ReadyMix (KAPA Biosystems, Wilmington, MA)], and amplified by PCR with the following conditions: 95°C for 3 min, [98°C for 20 sec, 65°C for 30 sec, 72°C for 5 min] x5, 72°C for 5 min, and hold at 4°C. The first WTA products were purified by an AMPure XP kit (Beckman Coulter, CA) at 0.7:1 ratio of beads to sample and eluted with 20 μL of 10 mM Tris-HCl pH 8.0. 10.5µL of the first PCR products were mixed with 14.5 μL of the second PCR mixture [0.4 μM of primers (5’ BDWTA V2 primer and illumina-i7-primer), and 1x KAPA Hifi Hotstart ReadyMix], and amplified with the same conditions as the first PCR. The second PCR products were purified by an AMPure XP kit at 0.7:1 ratio of beads to sample and eluted with 15 μL of 10 mM Tris-HCl pH 8.0, quantified by a Nanodrop 8000 (Thermo Fisher), and their size distribution was analyzed by a MultiNA system (Shimazu). 100 ng of the WTA library was subjected to fragmentation/end-repair/A-tailing using NEBNext Ultra II FS DNA Library Prep Kit for Illumina (New England Biolabs). Purified adapter-ligated products were subjected to index PCR. The PCR products were purified by double size-selection by using an AMPure XP kit at 0.5:1 ratio and 0.8:1 ratio (final 1.3X) of beads to sample and eluted with 12 μL of dH2O. Amplified products were analyzed by a MultiNA system. Final transcriptome libraries, with lengths of around 300 base pairs, were quantified using the KAPA Library Quantification Kit (KAPA Biosystems). Pooled libraries were sequenced by an Illumina Novaseq 6000 S4 flowcell (67 bp read 1 and 140 bp read 2) using NovaSeq 6000 S4 Reagent Kit v1.5. Oligo sequences used for this study are shown in Supplementary Table 4.

### Data Processing of Bulk TCR Sequencing Reads

Data processing of TCR-seq was performed as previously reported^15^. Briefly, reads were trimmed and filtered with Cutadapt-3.2^37^ and PRINSEQ-0.20.4^38^. Filtered reads were then aligned to a reference mouse TCR V/D/J sequence registered in the international ImMunoGeneTics (IMGT) information system with MiXCR-3.0.5^39^.

### Data Processing of Single Cell Sequencing Reads

Data processing of single-cell RNA-seq reads was performed as previously reported^15^. Resultant count data were converted to a single-cell gene-expression matrix file and filtered for valid cells based on read counts. Expression data of targeted panel genes were converted to a Seurat object^40^, log normalized and scaled (regressed by the number of unique molecular identifiers per cell) using the NormalizeData and ScaleData functions respectively. Principal component analysis (PCA) was performed using RunPCA function, and enrichment of each PC was calculated using the JackStraw and ScoreJackStraw function, and PCs that were significantly enriched (*P* ≤ 0.05) were selected for clustering and dimensional reduction analysis. Dimensional reduction was performed using RunUMAP function with default settings. Cell clustering was performed using FindNeighbors and FindClusters (resolution = 0.6) against the significant PCs. The downstream single cell analysis was conducted with Scanpy 1.9.1^41^, including the differential gene expression analysis (rank_genes_groups).

### Data Processing of Bulk RNA Sequencing Reads

Data processing of bulk RNA-seq data was performed as previously reported^42^. Raw data was then analyzed following standard protocols using the edgeR package^43^. Genes were filtered with filterByExpr, counts normalized with calcNormFactors, dispersion characteristics computed with estimateGLMCommonDisp, estimateGLMTrendedDisp and estimateGLMTagwiseDisp, before a negative binomial generalized log-linear model was fit with glmFit. Statistical testing between groups was performed with glmLRT to rank genes before pathway analysis with fGSEA^44^. We used the same gene signatures as those described in Gene Signatures Scores (Supplementary Table 3).

### Merging T Cell Clones Across Different Datasets

Within the bulk TCR-seq datasets, T cell clones were determined as TCR reads with the same TCRβ V segment, D segment, J segment, and CDR3 nucleotide sequence. Within the single-cell RNA/TCR-seq datasets, T cell clones were determined as cells with the same TCRβ CDR3 nucleotide sequence. Bulk TCR-seq datasets were merged based on shared TCRβ V segment, D segment, J segment, and CDR3 nucleotide sequences while bulk TCR-seq datasets and single-cell RNA/TCR-seq datasets were merged based on shared TCRβ CDR3 nucleotide sequences. Any cells in the single-cell RNA/TCR-seq datasets that shared TCRβ CDR3 nucleotide sequences with multiple clones in the bulk TCR-seq dataset of the same mouse were excluded from the single-cell RNA/TCR-seq analysis.

### Calculation of Clone Features

For the bulk TCR-seq experiments, clone frequency was calculated as the fraction of reads of a clone out of the total number of reads in the CD8^+^ T cell dataset for each mouse. For the single-cell RNA/TCR-seq and bulk TCR-seq experiments, clone frequency was calculated as the fraction of combined reads of a clone in all four bulk TCR-seq datasets, out of the total combined number of reads in all four datasets for each mouse. Read counts in each dataset were corrected by the total number of cells sorted to account for cells aliquoted for single-cell RNA/TCR-seq. Expansion of clones was calculated as the log2 fold-increase in clone frequency over the indicated days. Cell counts were calculated by correcting the bulk TCR-seq obtained frequencies with flow cytometry bead derived total CD8^+^ T cell counts. The fraction of cells PD-1^−^, precursor exhausted, intermediate exhausted and terminally exhausted for a clone was calculated by dividing the read count of the clone in each of the respective CD8^+^PD-1^−^, CD8^+^PD-1^+^Ly108^+^TIM-3^−^, CD8^+^PD-1^+^Ly108^+^TIM-3^+^ and CD8^+^PD-1^+^Ly108^−^TIM-3^+^ datasets by the combined read count of the clone across all four datasets. Clones (using their TCRβ CDR3 nucleotide sequence) were defined as expanding or contracting based on differential abundance analysis using the beta-binomial model trained on the default training dataset^22^. After identification of expanding and contracting clones (Fig. 1), subsequent analysis was restricted to clones that contained reads in the terminally exhausted (CD8^+^PD-1^+^Ly108^−^TIM-3^+^) dataset to enrich for tumor reactive clones^14^.

### Gene Signature Scores

Gene signature scores were calculated by normalizing and taking the log of the gene counts obtained from the pre-processing step and applying the score_genes function from Scanpy with the required gene signature, as in Tirosh et al., 2016^45^. Gene signatures from Im et al., 2016^46^, Beltra et al., 2020^47^, Galleti et al., 2020^48^, Liberzon et al., 2015^27^, Nath et al., 2019^31^ and Wells et al., 2023^30^ were used to score cells for a stem-cell like state, Tex^prog^^1^ and Tex^prog^^2^ cell states, T_PEX_ and T_SCM_/T_CM_ cell states, hallmark gene signatures, TGF-β signaling signature (restricted to genes with adjusted p-value<0.05 and fold-change>1) and Let-7 suppression signature respectively. Gene signatures are listed in Supplementary Table 3. Mouse ortholog genes^49^ (based on Ensembl Biomart version 87) were used when gene sets were derived from human data or needed to be converted for analysis on human data. Scores were produced for each cell, and these were averaged across all cell members of a clone to calculate mean expression scores.

### Analysis of Published Mouse Dataset

Single-cell RNA/TCR-seq dataset of antigen-signaled CD8^+^ T cells infiltrating the YUMMER1.7 tumor model^32^ (GSE266361) was downloaded and reprocessed as above using Scanpy with default parameters. Cells from the day 8 intraperitoneal dataset were subset from the data, log normalized, scored for gene signatures, merged with batch balanced kNN^50^ and clustered (resolution = 1.5) with new neighborhood coordinates. One cluster expressing Tcf7 and Lag3 in the tumor was identified as the precursor exhausted cell cluster. Clones were defined as cells with the same TCRα and β CDR3 nucleotide sequence.

### Analysis of Published Human Datasets

Single-cell RNA/TCR-seq data of site-matched tumors from Yost et al. 2019^13^ (GSE162498) was downloaded and reprocessed as above using Scanpy with default parameters. Cells from the CD8^+^ T cell clusters (as defined by the original authors: CD8_mem, Naive, CD8_act, CD8_ex, CD8_eff, CD8_ex_act) were subset from the data, log normalized and scored for the expansion gene signature. Analysis was restricted to CD8^+^ T cells from the ‘Exhausted’, and ‘Activated/ exhausted’ clusters (CD8_ex, CD8_ex_act) as the original authors described these clusters enriching for tumor-responding clones. Clone expansion analysis was restricted to clones from sample su009 as this was the only patient sample that contained clones with more than three cells in the relevant clusters pre-treatment. Clones were defined as cells with the same TCRα and β CDR3 nucleotide sequence. Patient responses were defined as described in the original publication. Bulk RNA-seq data of tumor biopsies from Riaz et al., 2017^5^ (GEO: GSE91061) and Gide et al., 2019^6^ (ENA: PRJEB23709) were processed and scored for the TCR.strong and expansion signature (Supplementary Table 2) as described in Elliot et al., 2021^33^. The Kaplan-Meier estimator of survival function was calculated with scikit-survival^51^.

### Software Versions

Data was analyzed using R version 4.0.3 and R packages (Seurat 4.0.4, edgeR 4.0.16, fGSEA 1.28.0), Python version 3.8.6 and Python packages (jupyterlab 4.1.5, numpy 1.24.4, pandas 1.5.2, scipy 1.10.1, scanpy 1.9.1, anndata 0.7.5, scikit-survival 0.22.2). Figures were produced with fGSEA 1.28.0 and EnhancedVolcano 1.20.0 in R, seaborn 0.11.0 and matplotlib 3.5.0 in Python, Prism 10, Affinity Publisher 1.10.8, and using illustrations from Irasutoya (https://www.irasutoya.com/).

### Statistical Analysis

Differential gene expression on the single-cell RNA/TCR-seq data was performed using Wilcoxon rank-sum test corrected with the Benjamini–Hochberg procedure and recorded for genes with p<0.05. Correlations were assessed using linear least-squares regression and two-tailed Wald Test. Distributions were assessed for significance using Kruskal-Wallis with Bonferroni correction where appropriate. 95% confidence intervals were calculated based on the data’s bootstrap distribution. For comparison of Kaplan Meier survival curves, patients with survival data were split into “High” and “Low” cohorts based on the on-treatment gene signature scores and analysed using a Log-rank (Mantel-Cox) test.

## Data Availability

Sequencing data that support the findings of this study have been deposited in the Gene Expression Omnibus database (GEO: available at time of publication). All other data supporting the findings of this study are available from the corresponding author on reasonable request.

## Code Availability

No new algorithms were developed for this manuscript. Code generated for display of each figure is deposited on Zenodo (available at time of publication). Additional code generated for analysis is available from the corresponding authors upon reasonable request.

## Acknowledgments

This work was supported by the Japan Society for the Promotion of Science under grants 17929397, 20281832, 22H05064, and 23H02706 and the Japan Agency for Medical Research and Development under grants JP22fk0310509, JP22fk0310514, JP22ama221306, and JP21gm6210025. J.E.D.T. was funded by the Medical Research Council (MC_UU_0025/12) award. D.B. was funded by the Medical Research Council (MR/V009052/1) and a Lister Institute of Preventive Medicine fellowship. M.T. was supported by the Masason Foundation, Recruit Foundation and the Gates Cambridge Trust. We thank Y. Hara for advice regarding cell sorting; members of IGT, Inc. for expert technical assistance with TCR sequencing; N. Yamaguchi, M. Nakano and Y. Kawamura for helpful discussions.

## Author contributions

M. Takahashi, M.Tsunoda, H.A. and S.U. conceived the project and designed the experiments. M. Tsunoda performed the experiments with help from H.A., M.K., H.O. and S.U. M. Takahashi analyzed the data with help from H.A., S.S., D.B. and M.Tsunoda. S.U. and K.M. supervised the project with support from J.E.D.T. and I.S. M.Takahashi, M.Tsunoda and S.U. wrote the manuscript. All authors reviewed the results and approved the final version of the manuscript.

## Materials & Correspondence

Correspondence and requests for materials should be addressed to Satoshi Ueha.

**Extended Data Fig. 1.**
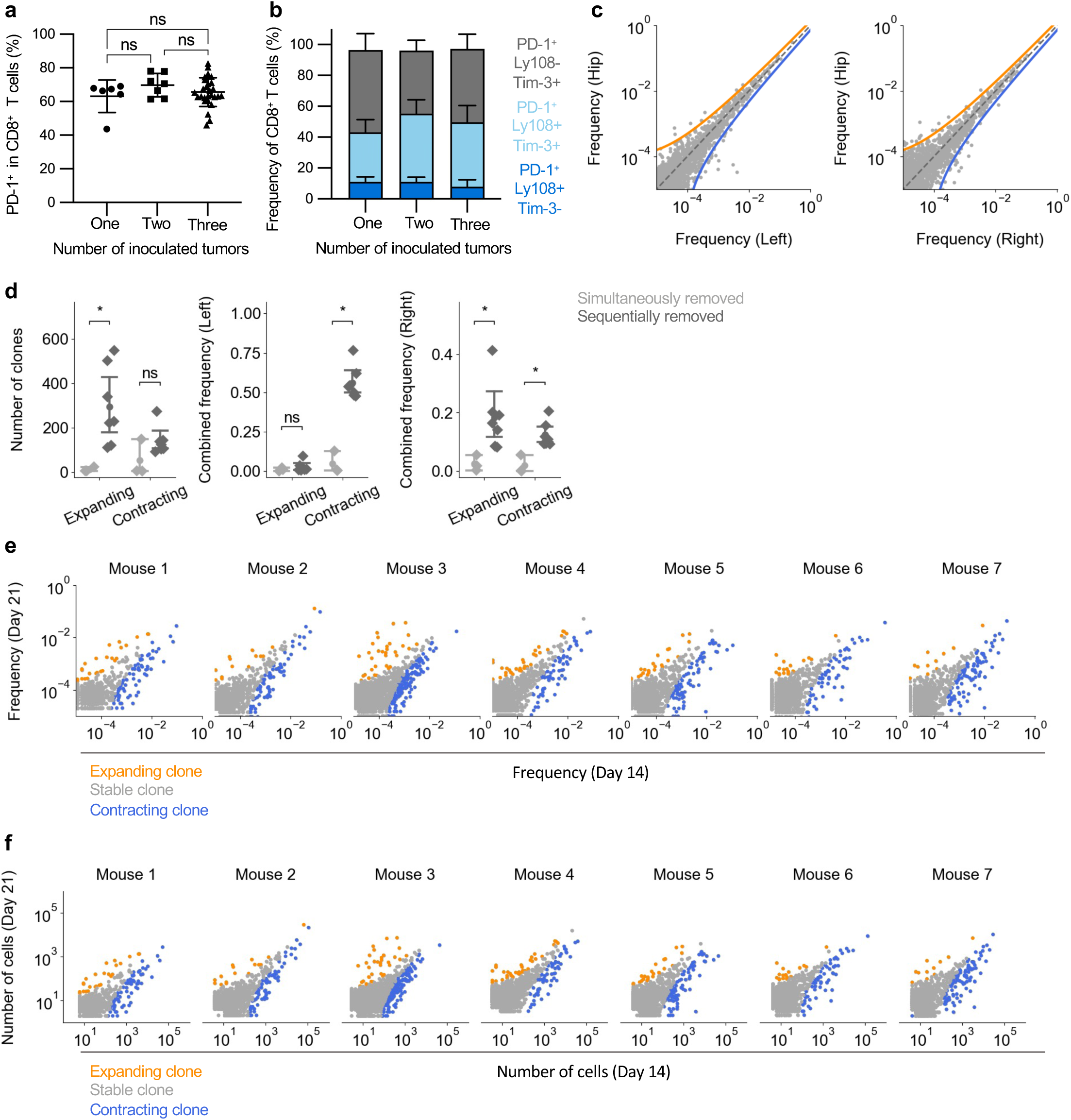
Multi-tumor mouse model reproducibly captures expanding and contracting clones – related to Fig. 1. **a-b)** Comparing the phenotypes of CD8^+^ T cells in tumors 14 days after mice were inoculated with different number of tumors. Fractions calculated by averaging across the tumors in each mouse. **a)** Fraction of PD-1^+^ cells in CD8^+^ T cells. **b)** Distribution of cell subsets (based on PD-1, Ly108 and TIM-3 markers) in CD8^+^ T cells. **c)** Scatter plots displaying the frequency of clones (normalized read count of each clone) in the left flank and left hip (Left) and right flank and left hip (Right) of 3 mice excised at the same time with representative bounds defining expanding and contracting clones overlayed. **d**) The number (Left), the combined frequency in the left tumor (Middle) and the combined frequency in the right tumor (Right) of expanding and contracting clones from tumors resected simultaneously (Light grey, n=3) and sequentially on days 14 and 21 (Dark grey, n=7). **e-f)** Scatter plots displaying the frequency (**e**) and cell counts (**f**) of clones in tumors on days 14 and 21 for individual mice. Clones are colored by their expansion dynamics. Dots represent mice (**a, d**) and clones (**c, e, f**) with mean and standard deviation (**a, b**) or 95% confidence interval as shown (**d**). Statistical testing via Kruskal-Wallis test with Bonferroni correction (**a, d**) (****, p<0.0001; ***, p<0.001; **, p<0.01; *, p<0.05; ns p>0.05).

**Extended Data Fig. 2.**
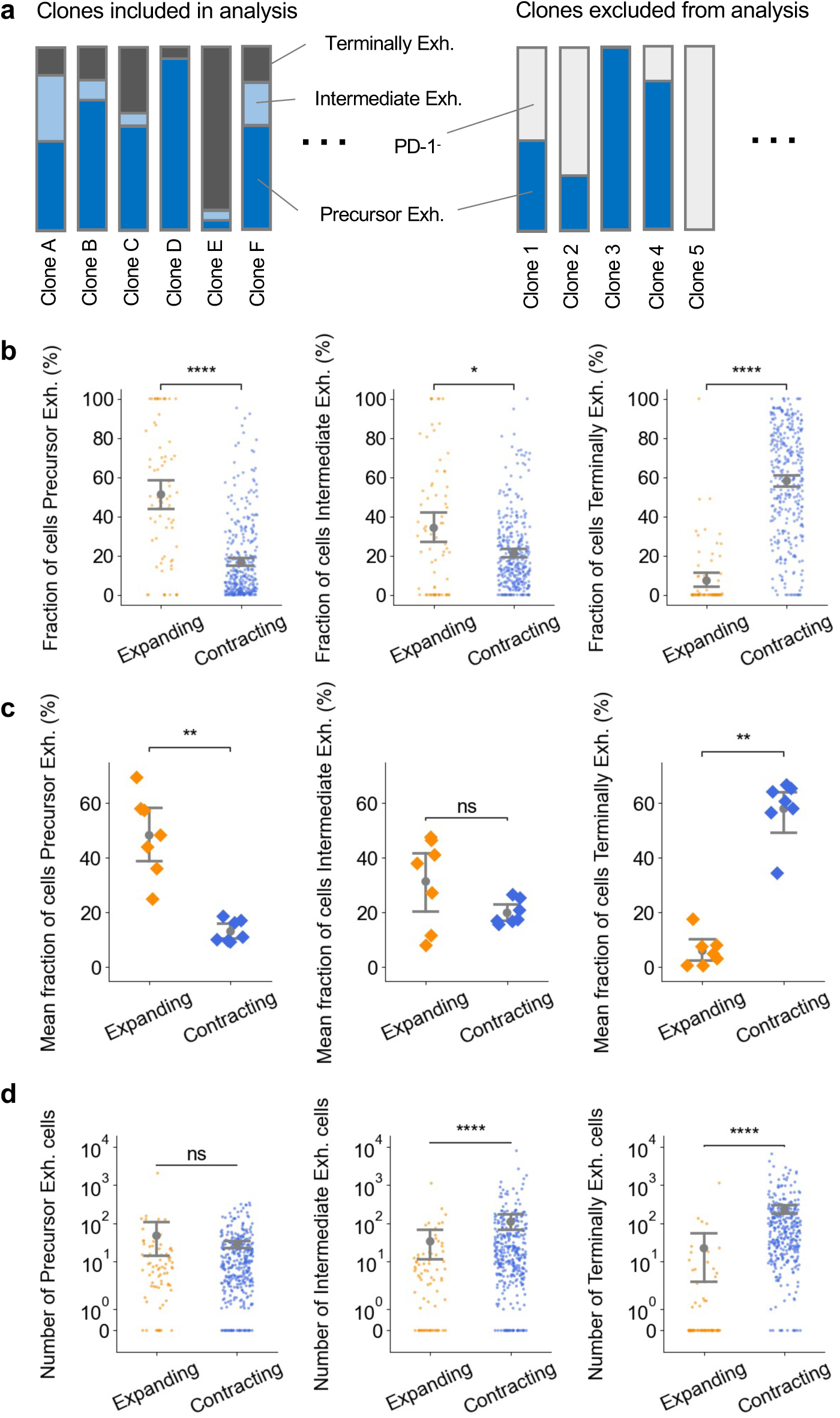
Expanding and contracting clones have distinct cellular compositions – related to Fig. 2. **a**) Diagram depicting filtering strategy used to enrich for tumor reactive clones. Merging of bulk TCR-seq datasets from sorted differentiation states enable the reconstruction of clones with their cellular compositions. Only clones with reads in the terminally exhausted dataset were selected for downstream analysis. **b-d**) Quantitative analysis of clones displayed in Fig. 2b. **b)** The fraction of precursor exhausted (Left), intermediate exhausted (Middle) and terminally exhausted (Right) cells of expanding and contracting clones. **c)** The mean fraction of precursor exhausted (Left), intermediate exhausted (Middle) and terminally exhausted (Right) cells of expanding and contracting clones in each mice. **d)** The number of precursor exhausted (Left), intermediate exhausted (Middle) and terminally exhausted (Right) cells of expanding and contracting clones. Dots represent clones (**b, d**) and mice (**c**) with mean and 95% confidence interval as shown (**b-d**). Statistical testing via Kruskal-Wallis test (**b-d**) (****, p<0.0001; ***, p<0.001; **, p<0.01; *, p<0.05; ns p>0.05).

**Extended Data Fig. 3.**
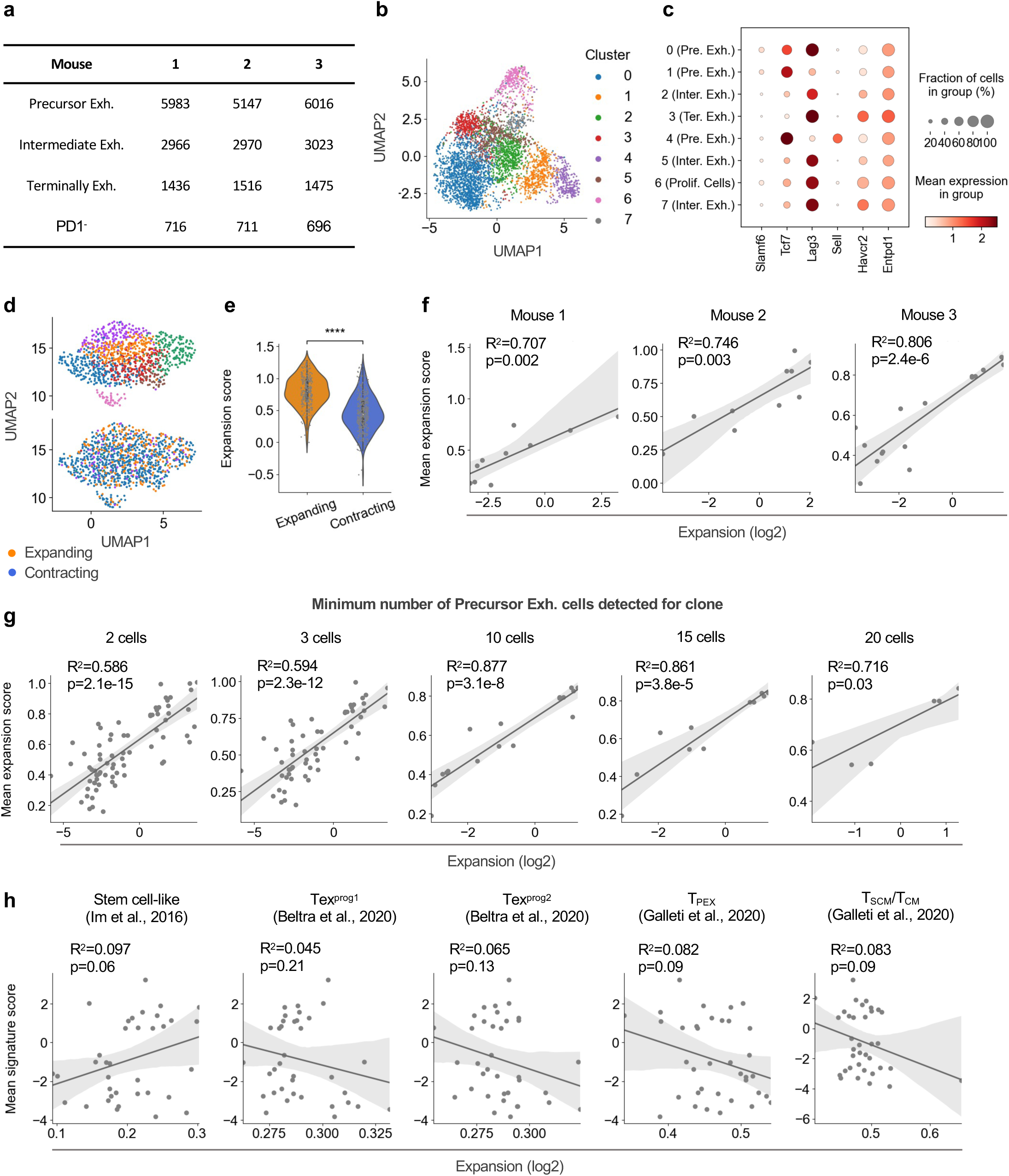
Expansion does not consistently correlate with previously reported precursor exhausted cell states – related to Fig. 3. **a)** Number of cells collected for each cell type from each mouse for single cell RNA/TCR-seq analysis on day 14. The remaining cells were processed through bulk TCR-seq. **b)** UMAP map of cells colored by clusters obtained from unsupervised Louvain clustering. **c)** Dot plot showing the expression of marker genes for each cluster. **d)** Re-calculated UMAP maps for precursor exhausted cells from expanding and contracting clones colored by clusters obtained from unsupervised Louvain re-clustering (Top) and clone expansion dynamics (Bottom). **e-g**) Cells (**e**) or clones (**f-g**) were scored for their expression of the expansion signature in precursor exhausted cells on day 14. **e)** Comparison of expansion signature score of cells from contracting and expanding clones. **f)** Scatter plots comparing the mean expansion signature score with the expansion (the log2 fold-increase in clone frequency from day 14 to 21) of clones for individual mice. Analysis on clones with at least 5 precursor exhausted cells on day 14. **g)** Scatter plots comparing the mean expansion signature score with the expansion of clones for different clone inclusion criteria. Inclusion criteria was altered based on the number of precursor exhausted cells in the single cell dataset for clones on day 14. **h)** Scatter plots comparing the mean signature score (of various precursor exhausted state gene signatures) with the expansion of clones. Analysis on clones with at least 5 precursor exhausted cells on day 14. Dots represent genes (**c**), cells (**b, d, e**) and clones (**f-h**). Statistical testing via Kruskal-Wallis test (**e**) and Wald test (**f-h**) (****, p<0.0001; ***, p<0.001; **, p<0.01; *, p<0.05; ns p>0.05).

**Extended Data Fig. 4.**
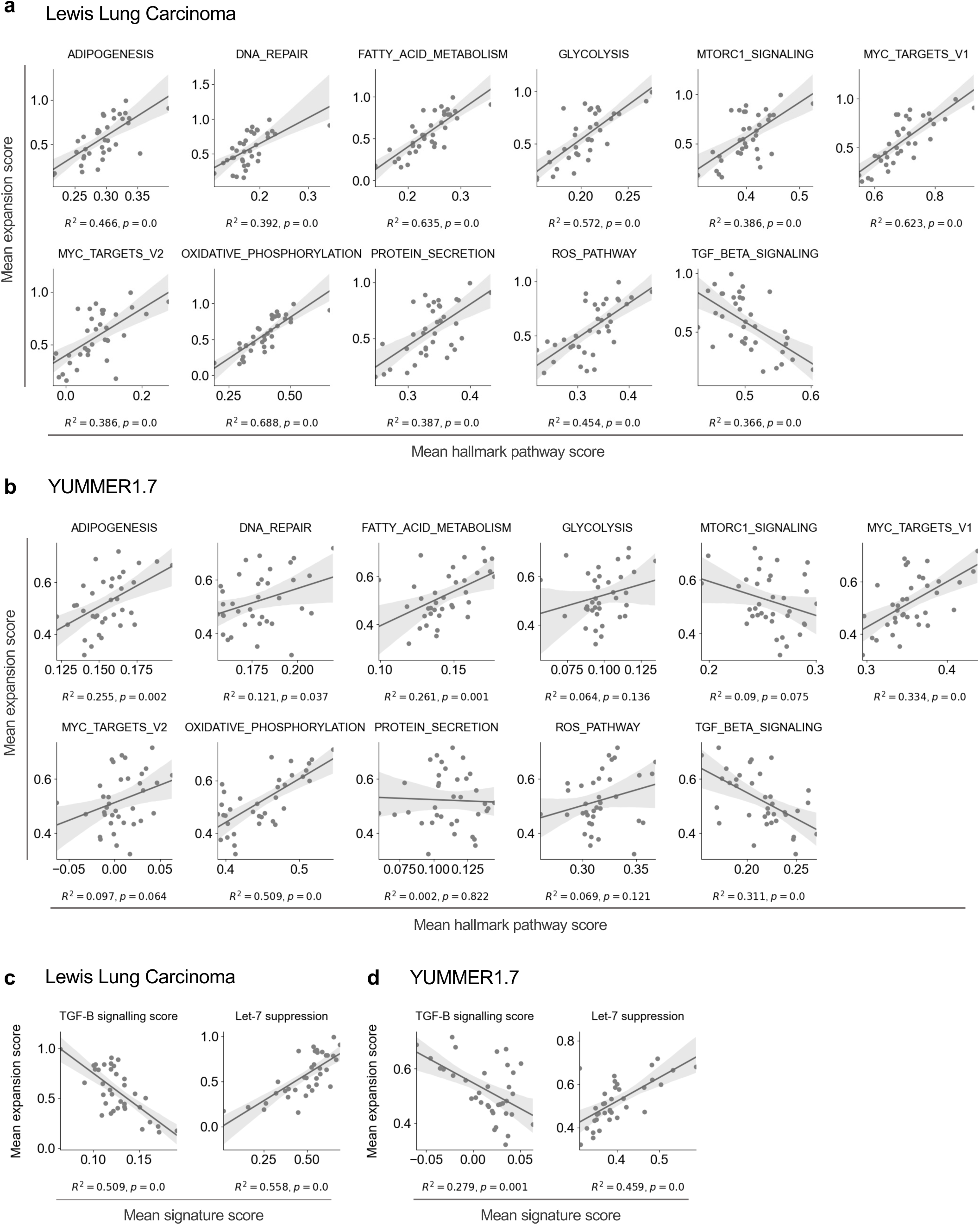
The expansion signature associates with high metabolic activity and reduced TGF-B signaling – related to Fig. 3. **a-b)** Scatter plots comparing the mean expansion signature score of clones with various mean hallmark pathway signatures for our Lewis Lung Carcinoma dataset (**a**) and a YUMMER1.7 single cell dataset from Takahashi et al., 2024^32^ (**b**). Analysis on clones with at least 5 precursor exhausted cells on day 14. Hallmark pathway signatures which strongly correlated (R^2^>0.3, p<0.05) with the mean expansion signature score in the Lewis Lung Carcinoma dataset are shown. **c-d)** Scatter plots comparing the mean expansion signature score of clones with a T cell TGF-B signaling score (Left) and Let-7 signaling suppression score (Right) for our Lewis Lung Carcinoma dataset (**c**) and a YUMMER1.7 single cell dataset (**d**). Analysis on clones with at least 5 precursor exhausted cells on day 14. Dots represent clones (**a-d**). Statistical testing via Wald test (**a-d**) (****, p<0.0001; ***, p<0.001; **, p<0.01; *, p<0.05; ns p>0.05).

**Extended Fig. 5.**
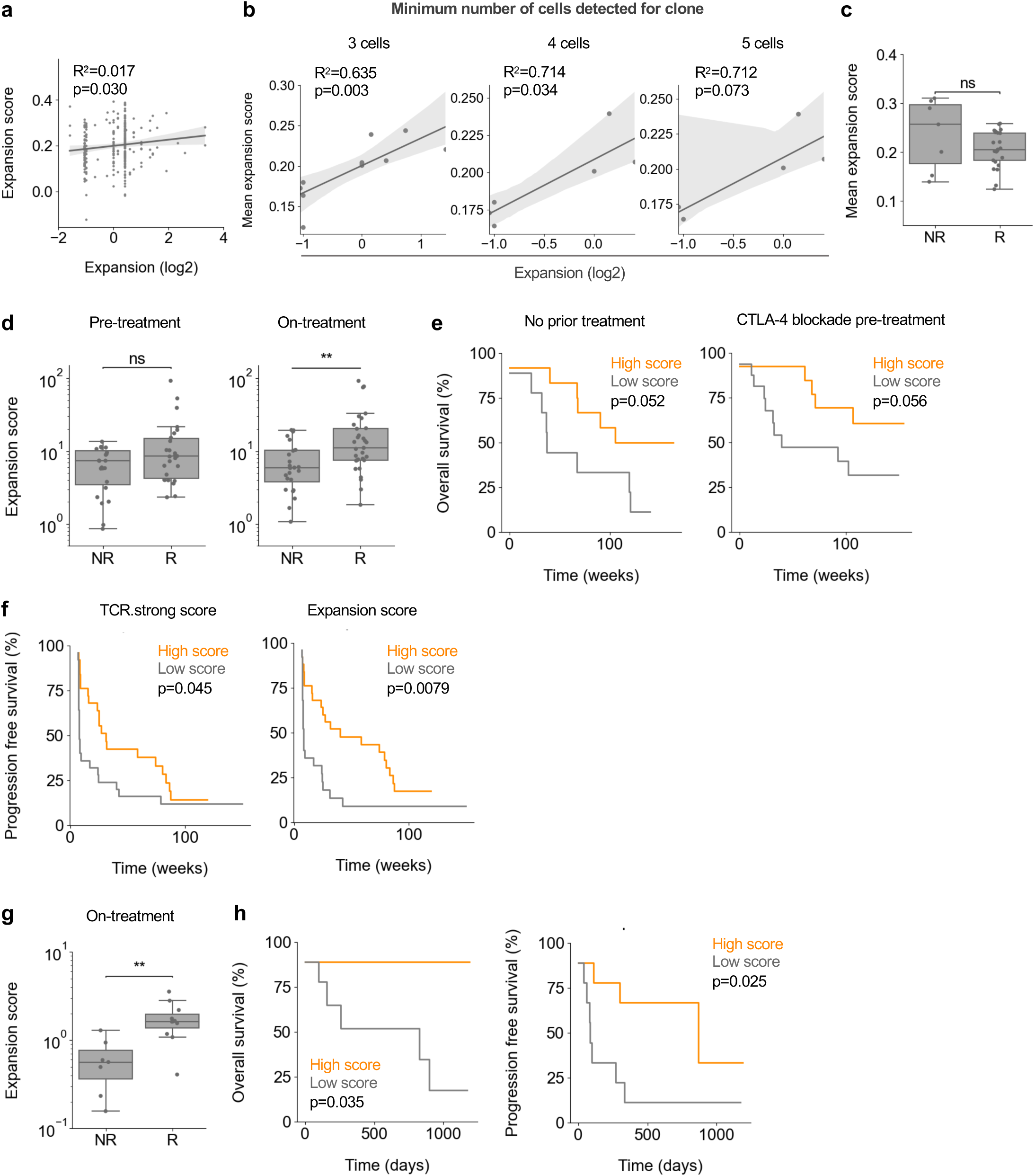
The expansion signature stratifies melanoma patient outcomes to different immunotherapy regimens – related to Fig. 4. **a-c)** Analysis of site-matched primary tumor single-cell RNA/TCR-seq data from Yost et al., 2019^13^. **a)** Scatter plot showing correlation between the expansion signature score of cells pre-treatment and the expansion of the clone (the log2 fold-increase of clone count from pre-to post-treatment). **b)** Scatter plots comparing the mean expansion signature score with the expansion of clones for different clone inclusion criteria. Inclusion criteria was altered based on the number of cells in the pre-treatment dataset for clones. **c**) Comparison of mean expansion score in cells of clones pre-treatment between non-responders (NR) and responders (R). Analysis of clones with at least 2 cells pre-treatment. **d-f)** Analysis of site-matched bulk RNA-seq tumor biopsies (n=50) from Riaz et al., 2017^5^. **d)** Comparison of expansion signature score in pre-treatment (Left) and on-treatment (Right) biopsies between non-responders (NR) and responders (R). **e)** Plots of Kaplan Meier estimator of overall survival based on median of expansion signature scores, splitting the cohort by their pre-treatment regimens: no prior treatment (Left, n=21) and CTLA-4 blockade pre-treatment (Right, n=29). **f)** Plots of Kaplan Meier estimator of progression free survival based on median of TCR.strong (Left) or expansion (Right) signature scores. **g-h)** Analysis of tumor biopsies (n=18) from Gide et al., 2019^6^. **g)** Comparison of expansion score in on-treatment biopsies between non-responders (NR) and responders (R). **h)** Plots of Kaplan Meier estimator of overall survival (Left) and progression free survival (Right) based on median of expansion signature score. Dots represent cells (**a**), clones (**b, c**) and patients (**d, g**). Statistical testing via Wald test (**a-b**), Kruskal-Wallis test (**c, d, g**) and Log-rank test (**e, f, h**) (****, p<0.0001; ***, p<0.001; **, p<0.01; *, p<0.05; ns p>0.05).

**Extended Data Fig. 6.**
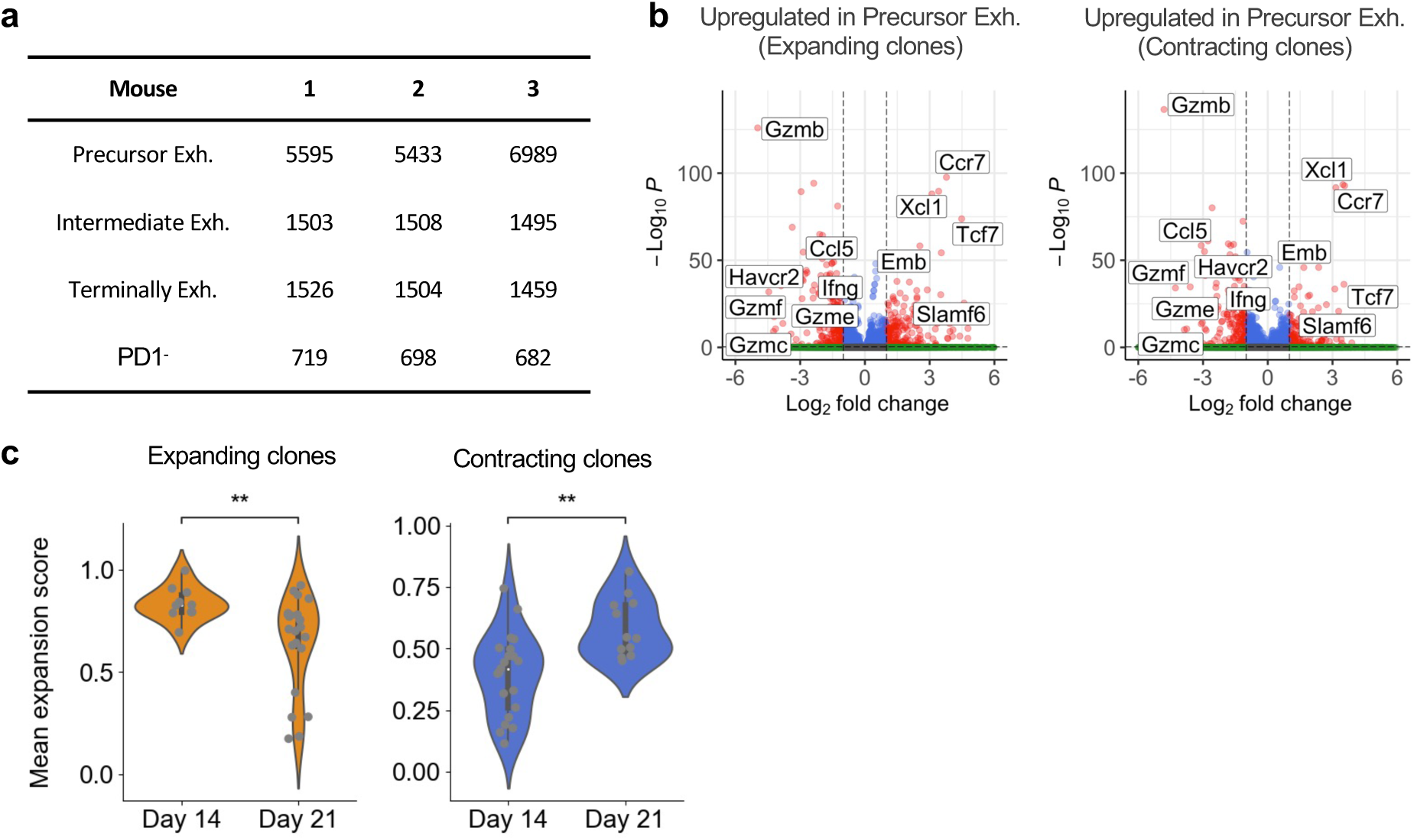
Precursor exhausted cells are maintained in the tumor during clone contraction – related to Figs. 2 and 3. **a)** Number of cells collected for each cell type from each mouse for single cell RNA/TCR-seq analysis on day 21. The remaining cells were processed through bulk RNA-seq. **b)** Differential gene expression analysis of genes overexpressed in precursor exhausted cells on day 21 from expanding (Left) and contracting (Right) clones over terminally exhausted cells. Notable genes are labelled. **c)** Change in mean expansion signature score of expanding (Left) and contracting (Right) clones between day 14 and 21. Analysis on clones with at least 5 precursor exhausted cells on day 14. Dots represent genes (**b**) and clones (**c**). Statistical testing via via Wilcoxon rank-sum test corrected with the Benjamini–Hochberg procedure (**b**) and Kruskal-Wallis test (**c**) (****, p<0.0001; ***, p<0.001; **, p<0.01; *, p<0.05; ns p>0.05).

**Extended Data Fig. 7.**
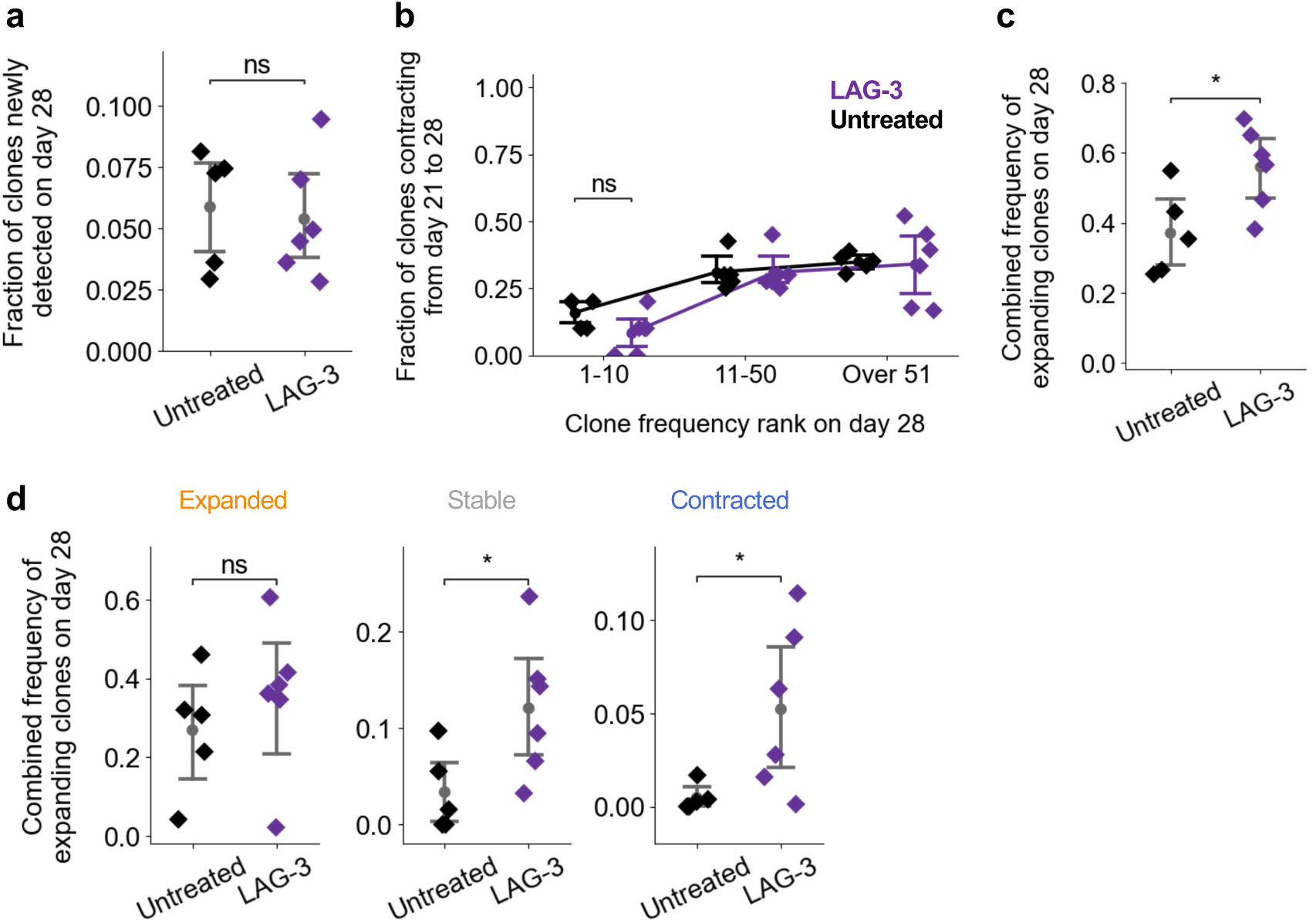
LAG-3 blockade increases the combined frequency of expanding clones – related to Fig. 5. **a)** Fraction of clones detected in the tumor on day 28 that were not detected in the tumors on days 14 and 21. **b)** Fraction of clones detected in the tumor on day 28 contracting between day 21 and 28, grouped by the ranked size of the clone on day 28 and colored by each treatment regimen. **c)** Combined frequency of clones on day 28 that expanded between day 21 and 28. **d)** Combined frequency of clones expanding between day 21 to 28 on day 28 for clones that expanded (Left), were stable (Middle) or contracted (Right) between day 14 and 21 (showing top 50 clones on day 14 or 21). Dots represent mice (**a-d**) with mean and 95% confidence interval as shown (**a-d**). Statistical testing via Kruskal-Wallis test (**a-d**) (****, p<0.0001; ***, p<0.001; **, p<0.01; *, p<0.05; ns p>0.05).

